# Duplicated binding site for RIC-3 in 5-Hydroxytryptamine Receptors Subtype 3

**DOI:** 10.1101/2022.02.17.480943

**Authors:** Hoa Quynh Do, Michaela Jansen

## Abstract

Serotonin or 5-hydroxytryptamine receptors type 3 (5-HT_3_) belong to the family of pentameric ligand-gated ion channels (pLGICs), which also includes other neurotransmitter-gated ion channels such as nicotinic acetylcholine receptors (nAChRs). pLGICs have been long-standing therapeutic targets for psychiatric disorders such as anxiety, schizophrenia, and addiction, and neurological diseases like Alzheimer’s and Parkinson’s disease. Due to structural conservation and significant sequence similarities of pLGICs’ extracellular and transmembrane domains across the more than 40 subunits found in humans, clinical trials for drug candidates targeting these two domains have been largely hampered by undesired effects mediated by off-subunit modulation. With the present study, we explore the interaction interface of the 5-HT_3A_ intracellular domain (ICD) with the resistance to inhibitors of choline esterase (RIC-3) protein. Previously, we have shown that RIC-3 directly interacts with the ICD of 5-HT_3A_ subunits. Using a sequential deletion approach, we identified the L1-MX segment of the ICD fused to maltose-binding protein as sufficient for the interaction. For the present study, synthetic L1-MX-based peptides, Ala-scanning, and a pull-down assay identified positions W347, R349, and L353 as critical for binding to RIC-3. In complementary studies with full-length 5-HT_3A_ subunits, the identified Ala substitutions reduced the modulation of functional surface expression by co-expression of RIC-3. Additionally, we found and characterized a duplication of the binding motif at the transition between the ICD MA-helix and transmembrane segment M4. Analogous Ala substitutions at W447, R449, and L454 disrupt MAM4-peptide RIC-3 interactions and reduce modulation of functional surface expression. In summary we identify two binding sites for RIC-3 with a shared duplicated motif in 5-HT_3A_ subunits, one in the MX-helix and one at the MAM4-helix transition.

## Main

Serotonin or 5-hydroxytryptamine type 3 receptors (5-HT_3_) are members of the family of pentameric ligand-gated ion channels (pLGICs) that also includes nicotinic acetylcholine receptors (nAChR)^1,2^. Gating of 5-HT_3_ channels by the endogenous agonist serotonin results in permeation by cations and signal transduction in the nervous system^3-10^. pLGICs’ involvement in brain-gut signaling circuitry and other non-serotonergic synaptic activities render these receptors effective and potential therapeutic targets for the treatment of conditions such as irritable bowel syndrome, chemotherapy-induced vomiting, inflammation, and psychiatric disorders^1,11-16^ 5-HT_3_ receptors assemble either into homopentamers from 5-HT_3A_ subunits or heteropentamers by combining 5-HT_3A_ subunits with other subunits (B through E)^17-19^. 5-HT_3A_ subunits share their molecular structure with the pLGIC family. Each homologous subunit comprises three domains: an extracellular N-terminal domain housing agonist-binding sites (ECD), a transmembrane domain enabling cations to cross the plasma membrane (TMD), and an intracellular domain playing critical roles in channel conductance and functional expression^20-22^ (ICD). The transmembrane domain consists of four α-helical transmembrane-spanning segments called M1, M2, M3, and M4 (Fig. 1a). In 5-HT_3A_ subunits, M3 and M4 are connected by a 115-amino acid segment that contributes the main part of the ICD. Structures of pLGICs solved by X-ray crystallography, cryo-EM, solution NMR, and EPR demonstrate that the ICDs of cation-conducting pLGICs consist of a short loop (L1) that connects M3 with a short helical segment (MX), and a large loop (L2) leading into the membrane associated (MA) and M4 helix (Fig.1)^21-24^.

**Fig. 1:**
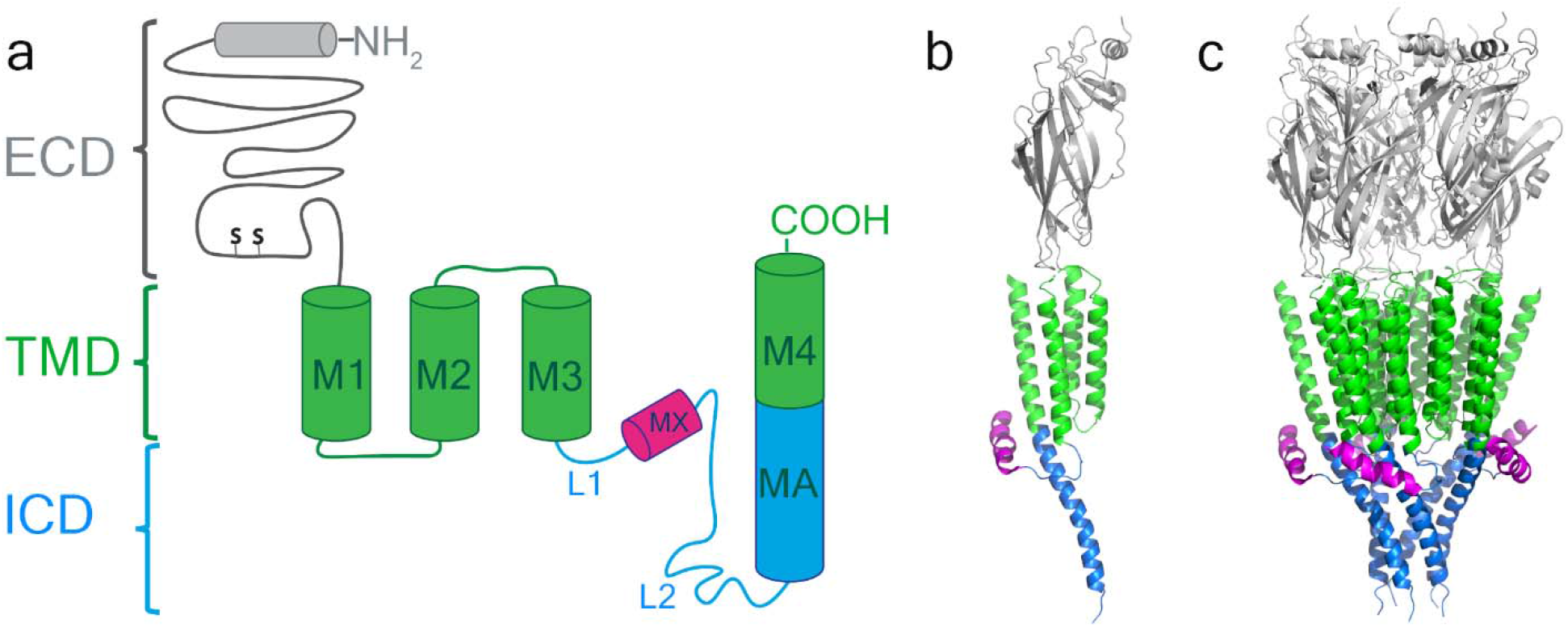
Domain structure of the 5-HT_3A_ receptor. **a**, Schematic of a single subunit illustrating the organization of the three domains, extracellular (gray), transmembrane (green) with four α-helical segments (M1 through M4), and intracellular (blue and magenta) with a 17 amino acid long MX-helix and a membrane associated (MA) helix of 31 amino acids in length continuous with the subsequent M4 transmembrane helix. **b**, and **c**, Ribbon representation of a single subunit and pentameric receptor (PDB: 4PIR)^1^. Colors as in a. All views parallel to the membrane plane.

Within the pLGIC superfamily of more than 40 subunits in humans, the M3-M4 cytoplasmic polypeptide chains or ICDs exhibit significant diversity in length, amino acid composition, and function. Originally, this domain had been considered largely disordered^21,22,25-28^. ICDs regulate gating and ionic conductance^29-31^, they mediate channel assembly and trafficking via interactions with intracellular proteins^23,32-34^. Of these intracellular proteins, the endoplasmic reticulum-resident Resistance to Inhibitors of Cholinesterase type 3 protein (RIC-3), was found to affect cell surface expression or functional expression of nicotinic acetylcholine (nAChR) and serotonin type 3A receptors (5-HT_3A_R) ^30,35-45^.

Expression of RIC-3 in *Xenopus leavis* oocytes enhances the whole-cell current amplitudes of human and rat α7 nAChR but reduces the current amplitudes of human α4β2 and α3β4 nAChRs ^35,40^, and also of 5-HT_3A_R^40,46,47^. In contrast, in mammalian cells, RIC-3 robustly enhances the cell surface expression of 5-HT_3A_ receptors and α4β2 and α3β4 nAChRs^36-38^. The direction of the RIC-3 effect additionally depends on its expression level. At low levels, RIC-3 was found to specifically regulate the assembly of α7 nAChR and to upregulate α4 and β2 subunit expression in a dose-dependent manner; however, at a high levels, RIC-3 inhibits assembly and cell-surface delivery of α7 nAChR; and this inhibitory action of RIC-3 is attributed to its self-aggregation that retains α7 in the endoplasmic reticulum^48-50^.

The extracellular (ECD) and transmembrane domains (TMD) of pentameric neurotransmitter-gate ion channels are long-standing therapeutic targets for psychiatric and neurological disorders. However, the highly homologous sequences and resulting conserved structures of ECDs and TMDs lead to interactions of drug candidates with off-target subunits, and consequently contributed to the discontinuation of many late clinical trials by Pfizer, Astra Zeneca, GlaxoSmithKline, Bristol-Mayer Squibb^21,22,26-28,51^.

To search for an effective subunit-specific approach to regulate the diverse functional properties of pLGICs, we have explored the interaction of the ICD of the 5-HT_3A_ receptor and RIC-3. Our previous studies on this interaction indicate a direct interaction between the ICD and RIC-3 that is mediated via the L1-MX segment of the ICD^47,52^. However, insights into the interaction sites or the molecular binding motif between RIC-3 and 5-HT_3A_ receptors remain elusive. A refined structural understanding of this interaction is expected to provide the basis for the structure-based design of new chemical entities (NCEs) interfering with this interaction and thus for the development of therapeutics for neurological and psychiatric disorders. Protein-protein interactions have been successfully employed in drug development^1,47,52-58^.

With the present study, using alanine-scanning mutagenesis, a protein-protein pull-down assay, and two-electrode voltage-clamp recordings, we provide evidence for the interaction between the 5-HT_3A_ L1-MX segment, as compared to the MBP-5-HT_3A_ L1-MX segment previously used^47,52,55^, and RIC-3 protein. We further demonstrate that further truncation to yield the MX-peptide is sufficient for the interaction with RIC-3. We identify a duplicated motif, DWLR…VLDR, present in both the MX-helix and at the transition of MA and M4-helices, essential for the interaction with RIC-3. Triple Ala substitution, W347, R349, and L353 in MX and W447, R449, and L454 in MAM4 disrupt RIC-3 interaction. In the 5-HT_3A_ structures, these motives are in close proximity at the interface between the lipid bilayer and cytosol.

## Results and Discussion

### The 5HT_3A_-MX helix alone interacts with RIC-3

In previous studies, we have demonstrated that RIC-3 directly interacts with the 5HT_3A_ receptor, specifically its ICD, and that the interaction is mediated via the L1-MX segment of the ICD^47,52,55^. In these prior studies, we used recombinant proteins fused with maltose-binding protein (MBP) to facilitate purification and increase stability of the isolated ICDs in the absence of the other domains as fusion partners. Both 6xHIS-MBP-RIC-3 and MBP-5-HT_3A_-ICD constructs in which segments of the ICD were sequentially deleted were expressed in *E. coli* and purified to homogeneity to explore protein-protein interaction. We demonstrated that the MBP-5-HT_3A_-L1-MX construct was the smallest construct investigated to mediate RIC-3 interaction. Here, we aimed at investigating the interaction of the 24-amino acid long 5-HT_3A_-L1-MX segment alone, without the highly-soluble and solubility enhancing MBP. To establish this modified system, we employed synthetic ICD peptides (Fig. 1a & Fig 2a&b) covalently linked to iodoacetyl resin and RIC-3 protein heterologously expressed in *Xenopus laevis* oocytes. The initial ICD peptides used were 5-HT_3A_-ICD-L1-MX with either a N- or C-terminal Cysteine (Cys). The Cys sulfhydryl was reacted with the iodoacetyl resin to form a stable thioether linkage. The peptide-resin was then used as the immobile phase together with solubilized RIC-3 protein in the mobile phase in a pull-down type assay. Non-covalently bound proteins were eluted with SDS-containing buffer, separated on SDS-PAGE and detected with RIC-3 antibody as described in detail in the method section. Representative data of RIC-3 expression in solubilized oocyte membranes and RIC-3 observed in eluates after pull-down using resin without and with L1-MX peptide covalently bound are shown in Fig. 2c.

**Fig. 2:**
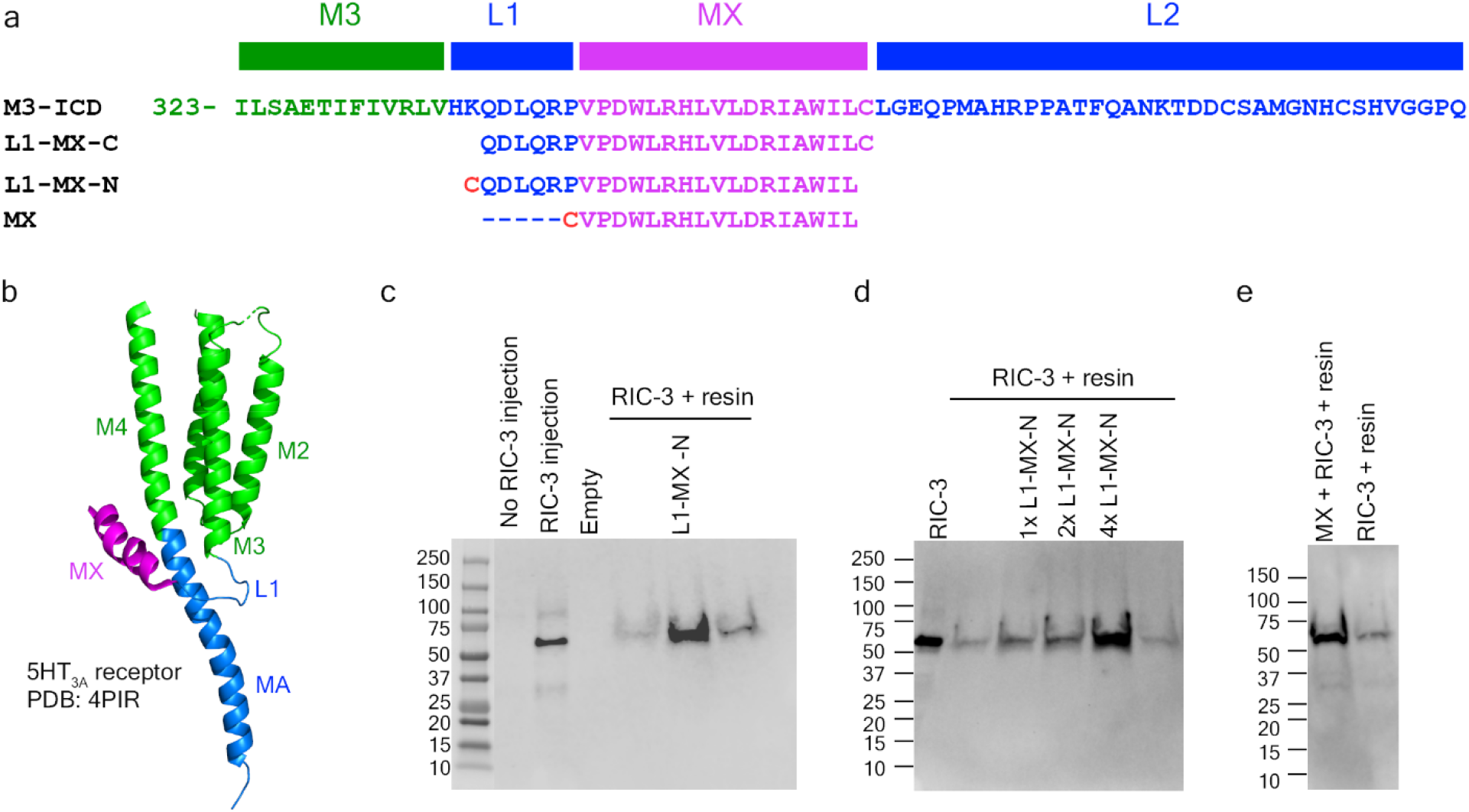
MX segment binds RIC-3. **a**, Sequence alignment of L1-MX- and MX-peptides with a partial 5-HT_3A_ sequence (top) consisting of the last 13 amino acids of M3 (green), L1-segment (blue), MX-helix (magenta) and 36 residues of the long L2-loop (blue) that connects with the MA-helix (blue, not shown in **a**, only in **b**). L1-MX-C contains a C-terminal Cys, and L1-MX-N and MX contain an N-terminal Cys. **b**, Ribbon representation of a single 5-HT_3A_ subunit indicating L1 and MX and other named peptide segments within the transmembrane and intracellular domains (PDB: 4PIR)^1^. **c** and **d**, Eluates of resin-linked peptide pull-downs of RIC-3. Western blot images show RIC-3 expressed in oocytes and control (no RIC-3 injection). Eluates from resin alone show the presence of a minor amount of RIC-3, whereas eluates from resin covalently linked with L1-MX-N peptide show prominent RIC-3 bands. 1x, 2x, and 4x indicate increasing peptide to resin ratios. **e**, Eluate from MX-linked resin yields RIC-3 band.

Without the injection of RIC-3 cRNA, no protein band was recognized by RIC-3 antibody in solubilized oocyte membrane samples (Fig. 2c). In contrast, a strong intensity band migrating just above 50 kDa was detected in the lane containing a sample from solubilized membranes of oocytes injected with RIC-3 cRNA. This result strongly indicates the presence of RIC-3 protein in solubilized oocyte membranes. Total membrane samples containing RIC-3 from three oocytes were then used for each pull-down assay reaction (Fig. 2c, rightmost lanes). Samples using resin without L1-MX peptide yielded a faint band indicative of weak/unspecific binding of RIC-3 to resin itself. In contrast, resin modified with L1-MX peptide at a peptide/resin ratio of 7.12 mg/ml, showed a dramatically increased band intensity. The result was consistently observed in at least ten independent pull-down experiments, and the data from 4 experiments are shown in Fig.2c & d. Based on that increasing band intensities were observed with increasing peptide to resin ratios (Fig. 2d), we used peptide to resin ratios above 4 mg/ml for subsequent experiments when utilizing 30 µl of settled resin to examine the interaction between RIC-3 and L1-MX peptides.

With the goal to precisely define the interaction interface between RIC-3 and 5HT_3A_ receptor, we then shortened the L1-MX peptide even further by removing the six amino acid long L1-segment (Fig. 2a). When using resin linked to the 17 amino acid MX-segment via an added N-terminal Cys for the RIC-3 pull-down, we obtained a much stronger band for RIC-3 as compared to the control sample with unlinked resin (Fig. 2e). We infer that the L1 loop does not play a critical role in the interaction between RIC-3 and the 5HT_3A_-ICD. The MX segment alone is sufficient for the RIC-3 and 5HT_3A_-ICD interaction. A previous study by Hales and co-workers found that when most of the amino acids in MX of the human 5-HT_3A_ receptor were removed in L-10(1) and L-10(2) constructs, the functional surface expression of human 5-HT_3A_ receptor in HEK cells was abolished^45^.

### Alanine substitutions at and between conserved sites, W347, R349, or L353, within the MX helix prevent interaction with RIC-3

To search for the RIC-3 binding interface within the 5-HT_3A_ MX segment, we used alanine scanning to generate single and multiple Ala replacements at conserved positions within the MX segment (Fig. 3a). These Ala substituted peptides were then immobilized on iodoacetyl resin as described above and used to probe RIC-3 binding. Single Ala substitutions W347A, R349A, or L353A in the MX segments gave rise to RIC-3 bands with a signal much stronger than that of the unspecific binding between resin and RIC-3. Under these conditions, the single Ala substitutions have little to no effect on the interaction between L1-MX and RIC-3 (Fig. 3b). However, the combined substitution of Ala residues for either seven amino acids from W347 to L353 (MX-7A peptide, Fig. 3a) or three conserved positions W347, R349, and L353 (MX-AAA peptide, Fig. 3a) in the MX segment drastically reduced the intensity of the RIC-3 bands to that of the unspecific or background binding between unmodified resin and RIC-3 (n=3). We infer that MX-7A and MX-AAA substitutions abolish the interaction with RIC-3 (Fig. 3c&d). Our data suggest that substitutions of conserved residues, W347, R349, and L353 in the MX segment, together, disrupt the interaction between ICD and RIC-3.

**Fig. 3:**
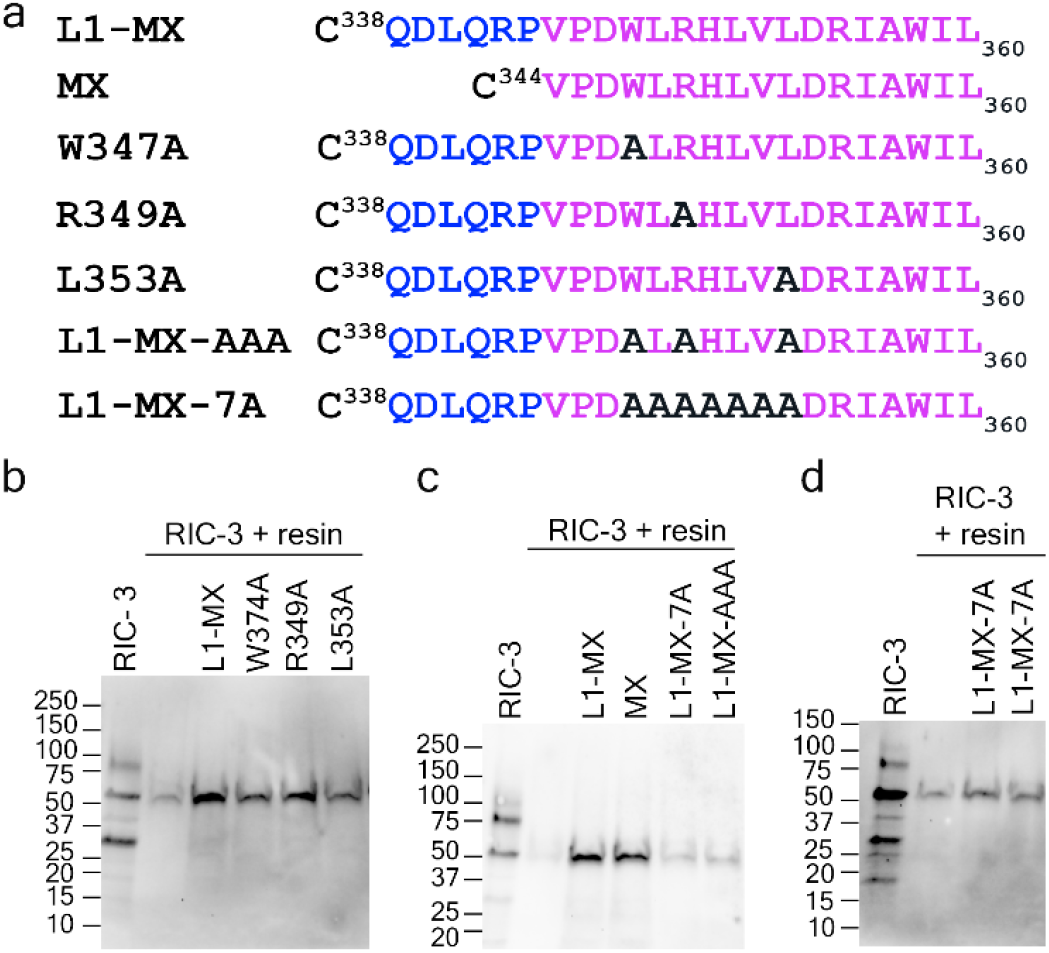
Alanine substitutions at conserved residues disrupt RIC-3 interaction. **a**, Sequence of L1-MX and MX peptides, and single and multiple site substitution peptides W347A, R349A, L353A, L1-MX-AAA, and L1-MX-7A. **b**, Single site substitution peptides W347A, R349A, or L353A in the MX helix preserve interaction with RIC-3, albeit with potentially altered affinity based on band intensities. **c** and **d**, Multiple Ala substitutions in L1-MX-AAA and L1-MX-7A disrupt the interaction with RIC-3. RIC-3 band intensities are comparable to Cys-capped resin background.

To address how the Ala substitutions at these conserved positions affect the function of the 5-HT_3A_ receptor and the modulation of its functional surface expression by RIC-3, we introduced these substitutions, W347A, R349A, and L353A, in full-length 5-HT_3A_ receptors and examined changes in the receptors’ function using *Xenopus* oocytes and two-electrode voltage-clamp recordings.

### Impact of Ala substitutions at W347, R349, and L353 on RIC-3 modulation of functional surface expression of 5-HT_3A_ receptors

We generated four constructs of 5-HT_3A_ receptors, i.e., three single Ala substitutions (W317A, R349A, or L353A) and one triple substitution (W317A, R349A, and L353A) (Fig. 4a).Complementary RNA (cRNA) for these engineered constructs and the wild type 5-HT_3A_ receptor were injected into *Xenopus* oocytes, either alone or with cRNA for RIC-3, at a ratio of 6 ng to 1.5 ng as examined previously^47^. Three days post-injection, the oocytes were used to examine the inhibitory effect of RIC-3 on the functional surface expression of 5-HT_3A_ channels by eliciting whole-cell currents with 10 μM of serotonin (5-HT). At least six oocytes from two to four different batches of oocytes were used for each type of 5-HT_3A_ receptor. Without the injection of RIC-3 cRNA (Fig.4b), no significant differences in current amplitudes were observed between the wild type channels (WT) and 5-HT_3A_ channels with a single Ala substitution, W347A, R349A, or L353A. However, the current amplitude elicited from 5-HT_3A_ channels containing the MX-AAA substitution was reduced to 47.7%±0.1 compared to WT (one-way ANOVA with Tukey’s post hoc test, p ≤ 0.0001). Our data suggests that single Ala substitutions W347A, R349A, or L353A do not interfere with the maximum current amplitude of the 5-HT_3A_ channels, whereas the triple substitution W347A, R349A, and L353A significantly reduces current amplitudes compared to WT. Our experiments cannot distinguish whether the inhibitory effect is mediated via a reduction in surface expression itself or an impact on function (e.g. open probability, single-channel conductance). To the best of our knowledge, this is the first time the functional role of residues W347A, R349A, and L353A has been reported. Previous studies on the M3-M4 loop or ICD of 5-HT_3A_ receptors or acetylcholine receptors from other groups did not specifically note a functional significance of W347, R349, and L353 residues^1,29-32,45,59,60^.

**Fig. 4:**
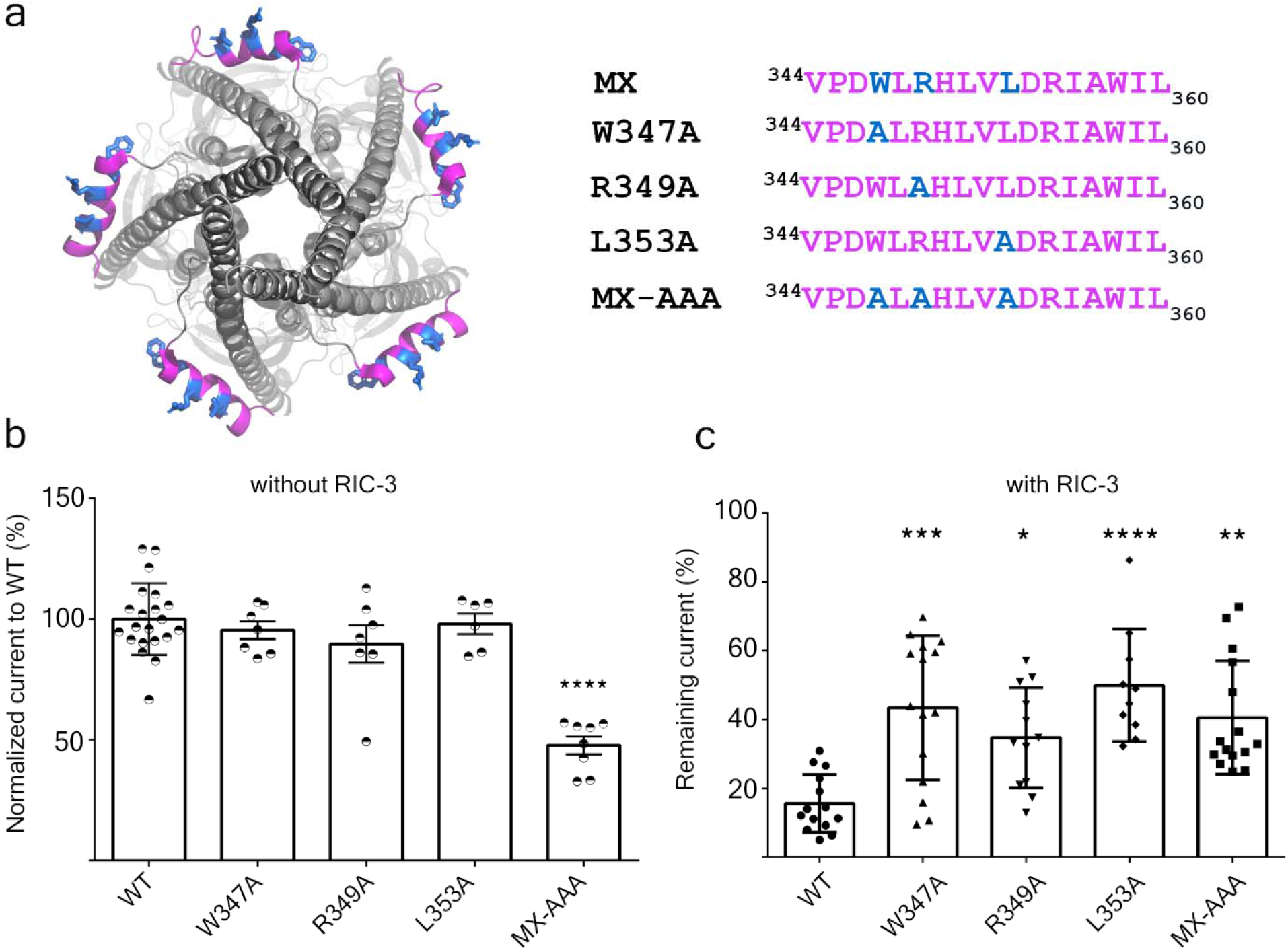
Ala substitutions within the MX-segment reduce modulation of functional 5-HT_3A_ surface expression by RIC-3. **a**, View of 5-HT_3A_ channel (PDB: 6W1J)^2^ from the intracellular side, with residues W347, R349, and L353 in blue (left), and alignment of the MX-segments with Ala substitutions as indicated (right). **b**, Serotonin-induced current amplitudes of all constructs normalized to the current amplitude of wildtype (WT) 5-HT_3A_ receptor. The triple Ala substitution leads to significantly reduced current amplitudes. **c**, Current amplitudes for each construct co-expressed with RIC-3 normalized to the amplitudes of each channel expressed alone, respectively. Statistical significance in **b** and **c** was determined vs. WT using ordinary one-way ANOVA with Tukey’s post-hoc test. Significance is indicated: * p ≤ 0.05, ** p ≤ 0.01, *** p ≤ 0.001, **** p ≤ 0.0001.

When 5-HT_3A_ channels were co-injected with RIC-3, we observed that RIC-3 inhibits 5-HT_3A_ current amplitudes elicited by serotonin in *Xenopus* oocyte as reported in previous studies^40,46,47^. The RIC-3 inhibitory effect was calculated as “remaining current” for each pair of 5-HT_3A_ receptors as follows: remaining current (%) = the peak current amplitude of 5-HT_3A_ channels in the presence of RIC-3 divided by the peak current amplitude from 5-HT_3A_ channels alone. This remaining current is used to assess the interaction between RIC-3 and the respective 5-HT_3A_ wildtype or modified receptors. For wild type 5-HT_3A_ channels, co-expression with RIC-3 lead to remaining currents of 15.62%±0.08. The remaining current of MX-AAA 5-HT_3A_ channels, 40.59%±0.16, was 2.6 times higher than that in wild type 5-HT_3A_ channels (one-way ANOVA with Tukey’s posttest, p < 0.01) (Fig. 4c). This result indicates that Ala replacements at residues

W347, R349, and L353 weakened the inhibitory effect of RIC-3 on the functional surface expression of 5-HT_3A_ channels. Similarly, the remaining currents for the single substitutions W347A, R349A, or L353A when co-expressed with RIC-3 showed a significantly reduced impact of RIC-3 co-expression compared to wild type. Significant changes in the inhibitory action of RIC-3, thus, were observed in all constructs containing Ala substitutions in the MX segment compared to wild type channels. RIC-3 co-expression reduced the currents to ∼50%, on average, in the single and triple substitutions compared to ∼16% in the wild type channels.

Overall, the full-length 5-HT_3A_ RIC-3 interaction studies corroborate that W347, R349, and L353 are important for RIC-3 interaction. However, the MX-AAA substitution’s impact on the interaction observed in the pull-down assay using an isolated 5-HT_3A_ segment and the in-vivo assay using full-length 5-HT_3A_ subunits is not equivalent quantitatively: the interaction was abolished in pull-down assay as compared to attenuated in the functional surface expression assay.

This difference in interactions of RIC-3 with either L1-MX peptides containing Ala substitutions or full-length 5-HT_3A_ mutants lead us to question if there is more than one binding site for RIC-3 within the 5-HT_3A_ subunit. We identified a duplication of the highly conserved MX-helix WRL motif (D**W**L**R**…V**L**DR) at the transition between MA and M4 helices (Fig. 5a). In the MA-M4 transition the corresponding residues are W447, R449 and L454.

**Fig. 5:**
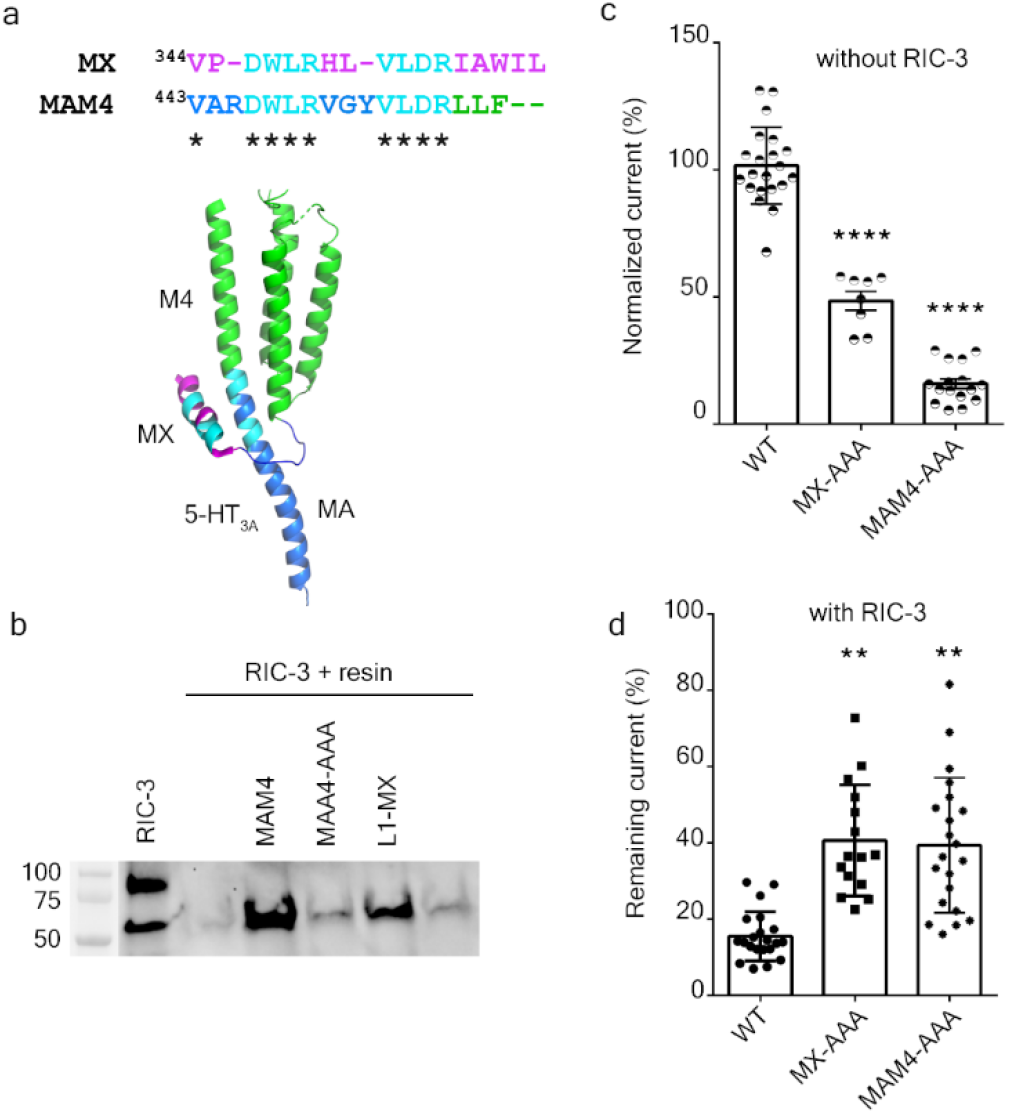
A duplicated RIC-3 binding motif at the MA-M4 transition mirrors the effects observed for the MX-segment RIC-3 site. **a**, Duplicated RIC-3 binding motif, DWLRxxxVLDR, depicted on the cartoon structure of 5-HT_3A_ transmembrane and intracellular domains: two amino acid clusters (cyan color) consisting of Asp, Trp, Leu, Arg (DWLR) and Val, Leu, Asp, Arg (VLDR), respectively, are present in both MX-helix and straddling the MA-M4 helices. **b**, Triple Ala substitution at residues W447, R449, and L454 of the MAM4 peptide reduces the RIC-3 band intensity to background levels. **c**, Serotonin-induced current amplitudes for AAA substitutions normalized to wildtype (WT) currents show significantly reduced amplitudes as compared to WT 5-HT_3A_ receptor. **d**, Remaining 5-HT current for each construct in the presence of RIC-3. AAA substitutions lead to less decreased currents with RIC-3 co-expression as compared to wildtype (WT). Statistical significance in **c**, and **d** was determined vs. wildtype (WT) using ordinary one-way ANOVA with Tukey’s post-hoc test. Significance is indicated: ** p ≤ 0.01 and **** p ≤ 0.0001.

### Duplicated RIC-3 motif at the MA-M4 transition

To test whether the duplicated motif in the MA-M4 transition presents a second binding site for RIC-3 within the 5-HT_3A_ subunit we therefore generated peptides and constructs containing Ala substitutions for residues W447, R449, and L454 and examined their impact using both the pull-down assay and electrophysiology to determine functional surface expression (Fig. 5b). Using L1-MX peptide and peptide-unmodified resin as the controls, we observed a strong RIC-3 signal in the pull-down assay using MAM4 peptide, which was almost abolished for MAM4-AAA. The Ala replacements at residues W447, R449, and L454, thus, disrupted the interaction between RIC-3 and MAM4 peptide similarly to the Ala replacements in L1-MX peptide (Fig. 3c).

In the absence of RIC-3 co-expression, full-length 5-HT_3A_ channels containing the triple substitution MAM4-AAA produced serotonin-induced currents with the amplitude equivalent to 15.65%±0.08 of the currents elicited for wild type, which is lower than the currents for the MX-AAA construct, 47.7%±0.1 (Fig. 5c). This data reveals that W447, R449, and L454 residues play a critical role in the functional expression of 5-HT_3A_ channels (WT vs. MAM4-AAA, one-way ANOVA with Tukey’s posttest, p < 0.0001). Residues W447, R449, and L454 also possess a stronger inhibitory effect on the 5-HT_3A_ functional expression than residues W347, R349, and L353 (one-way ANOVA with Tukey’s posttest, p < 0.0001).

When being co-expressed with RIC-3, the full-length 5-HT_3A_ receptors containing W447A, R449A, and L454A in the MA-M4 transition produced a remaining current of 36.88% ±0.18 (Fig. 5d). This inhibitory effect of RIC-3 is similar to the one determined for MX-AAA 5-HT3A channels with 40.59% ±0.16 seen in (Fig. 4c & 5d). Both of these remaining currents for MX-AAA and MAM4-AAA are significantly higher than the ones observed for wild type channels, 15.62%±0.08, as reported above (Fig. 4c and Fig. 5d).

Our findings thus confirm a second binding site for RIC-3 at the MA-M4 transition of 5-HT_3A_ ICD in addition to the site we identified within MX. Removing both RIC-3 sites in 5-HT3A would be expected to completely eliminate the effect of RIC-3 co-expression to reduce functional surface expression. The construct with six substituted positions (W347A, R349A, L353A W447A, R449A, and L454A) did not yield serotonin-induced currents either in the absence or presence of RIC-3.

The relative spatial orientation of both sites is visualized in Fig. 6. Intriguingly, the two duplicated motifs are in close proximity with the involved helices almost packing against each other but the key sidechains accessible on the surface. The sidechains of the conserved residues Trp, Arg, and Leu (WRL) in MX (Fig. 6a) point outward to the surrounding environment (close to the headgroup region of the lipid bilayer) in an orientation that primes these residues for participation in an interaction with a 5-HT_3A_ binding partner like RIC-3 that is also a transmembrane protein. This outward orientation of the WRL sidechains is observed in the majority of available structures of drug-bound or resting state 5-HT_3A_ receptors (PDB IDs: 4PIR, 6BE1, 6DG7, 6DG8, 6NP0, 6W1J). Residues Trp, Arg, and Leu in MX are conserved across species and subunits of cation-conducting pentameric ligand-gated ion channels. The corresponding residues in MX of nAChR α7 subunits (W330, R332, and L336, Uniprot code of P36544), also face outward to the surrounding environment (PDB ID 7KOQ). Previous studies demonstrate that RIC-3 enhances the functional expression of nAChR α7^43,61,62^. Of note, in many expression hosts there is only negligible nAChR α7 functional surface expression in the absence of endogenous or exogenous RIC-3 expression. nAChR α7 subunits only contain the conserved MX RIC-3 binding motif but not the duplicated motif in MAM4. The opposing effects of RIC-3 co-expression on nAChR α7 (increased expression) and 5-HT3A-MAM4-AAA (decreased expression) when only the MX site is present, indicate that the directionality of the effect depends on factors other than the pLGIC subunit RIC-3 interaction interface itself. Clearly, more studies are needed to dissect the molecular mechanism of RIC-3’s modulation of pLGIC functional surface expression.

**Fig. 6:**
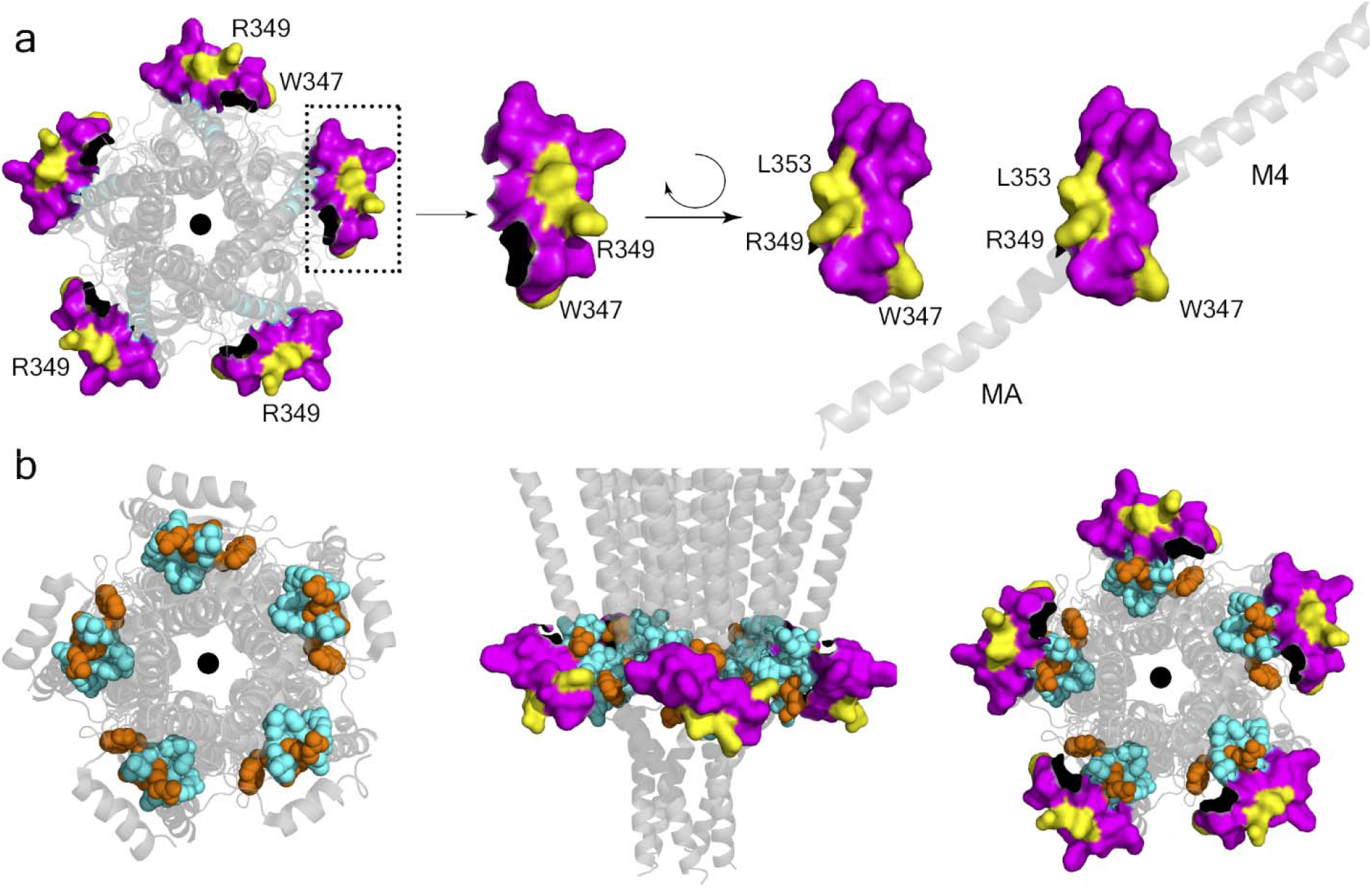
Visualization of RIC-3 binding motifs in the MX segment and in the MA-M4 transition. **a**, intracellular view of the pentameric 5-HT_3A_ structure. MX-helix is in magenta space filling representation, and amino acids W347, R349, and L353 of MX are in yellow. **b**, intracellular view with the MA-M4 transition in cyan and orange, and residues W447, R449, and L454 of MA-M4 are in orange, left; view parallel to the membrane in addition with MX segment in magenta and yellow as in **a**, middle; and intracellular view with the same color representation, right (PDB: 6W1J) ^2^.

## Conclusion

Here we identify a sequence motif in 5-HT_3A_ channels within the intracellular domain MX segment that mediates interaction with RIC-3. The motif is conserved across species and only observed in cation-conducting family members of pGLICs. Anion conducting family members do not contain this motif and their functional surface expression is also not modulated by RIC-3 co-expression. Intriguingly, we observe a duplication of this MX-helix binding motif DWLR…VLDR for RIC-3 at the MAM4 transition that is unique for 5-HT_3A_ subunits, but not other subunits. Within the completely folded pentameric 5-HT_3A_ channel, these two motifs are in close proximity. Our findings provide detailed insights into the interaction between the intracellular domain of pGLICs and RIC-3 that can be used to guide drug design aiming at regulating the functional expression of pLGICs and to further guide mechanistic studies of RIC-3 pLGIC modulation.

## Methods

### Mutagenesis

Single amino acid substitutions in the serotonin type 3A (5-HT_3A_) subunit, W347A, R349A, and L353A and two triple substitutions W347A, R349A, and L353A (or WRLAAA) and W447A, R449A, and L454A (or MA4-WRLAAA) were generated using appropriate primers and the QuikChange II XL Site-Directed Mutagenesis kit (Agilent Technologies) and were confirmed by DNA sequencing. The forward primers for the individual substitutions W347A, R349A, and L353A were 5’-CGGCCAGTACCTGAC**<u>GCA</u>**CTGAGGCACCTGGTC-3’, 5’-CAGTACCT-GACTGGCTG**<u>GCA</u>**CACCTGGTCCTAGACAG-3’, and 5’-CTGAGGCACCTGGTC**<u>GCA</u>**GA-CAGAATAGCCTGG-3’, respectively. The forward primer for the triple substitution WRLAAA was 5’-CAGTACCTGAC**<u>GCA</u>**CTG**<u>GCA</u>**CACC-TGGTC**<u>GCA</u>**GACAGAATAGCCTGGA-3’. The forward primer for the triple substitution MA4-WRLAAA was 5’-G**<u>GCA</u>**GTGGGATACGTG**<u>GCA</u>**GACAGGCTGCTG**<u>CTC</u>**C-3’. All DNA sequences were confirmed by Genewiz/Azenta.

### Complementary RNA (cRNA) production and oocyte injection

Expression plasmid vectors, pGEMHE19 and pXOON, carrying the human RIC3 sequence (GenBank: NM_001206671.2) and mouse 5-HT_3A_ sequence (GenBank: AY605711.1) with V5 epitope tag (GKPIPNPLLGLDSTQ) were generously provided by Dr. Millet Treinin (Hebrew University, Israel) and Dr. Myles Akabas (Albert Einstein College of Medicine, New York), respectively. Expression plasmid vectors containing substitutions of the mouse 5-HT_3A_ sequence were generated as described in the previous section.

The plasmid vector, 30 μg each, was linearized for 18 hours at 37°C in a 250 μl reaction containing restriction enzyme *Nhe*I and CutSmart buffer (New England Biolabs). The linear DNA plasmid was next used to produce complementary RNA (cRNA) of RIC-3 and 5-HT_3A_ wild type and engineered receptors. This *in vitro* transcription process was performed by employing the mMESSAGE mMACHINE T7 Kit (Ambion by Life Technologies). The cRNA was then purified using the MEGAclear kit (Ambion by Life Technologies), dissolved in nuclease-free water, and stored at -80°C until use.

### Expression of proteins in *Xenopus laevis* oocytes

Defolliculated *Xenopus laevis* oocytes were purchased from Ecocyte Bioscience US LLC (Austin, TX). Oocytes were transferred to standard oocyte saline (SOS) medium (*100 mM NaCl, 2 mM KCl, 1 mM MgCl*_*2*_, *1*.*8 mM CaCl*_*2*_, *5 mM HEPES; pH 7*.*5*) supplemented with 1x antibiotic-antimycotic (ThermoFisher Scientific/Gibco Cat. # 15240-062) and 5% horse serum (Sigma-Aldrich, St. Louis, MO, USA). Oocytes were injected on the same day of the delivery.To express RIC-3, 10 ng of cRNA (50 μl at 200 ng / μl cRNA) or 50 μl of sterile water as control were injected into each oocyte; the oocytes were then incubated at 15°C for 65-68 hours before being harvested for plasma membrane isolation. To co-express wild type (wt) or engineered (“mutant”, mt) 5-HT_3A_ receptors, with RIC-3, 6 ng of 5-HT_3A_-wt or 5-HT_3A_-mt cRNA and 1.5 ng of RIC-3 cRNA were injected into each oocyte, respectively. Oocytes injected with 5-HT_3A_-wt or 5-HT_3A_-mt cRNA alone were used as controls. After the injection, the oocytes were incubated at 15°C for 3 days before they were used for two-electrode voltage-clamp (TEVC) recordings. SOS medium was replaced 12 hours after the injection and subsequently every 24 hours.

### Isolation and solubilization of oocyte membranes containing RIC3

To isolate oocyte plasma membranes, oocytes were homogenized using a glass Teflon homogenizer and lysis buffer (*25 mM MOPS, 1 mM EDTA, 0*.*02 % NaN*_*3*_; *pH 7*.*5)* supplemented with protease inhibitor cocktail III (Research Products International, Mt. Prospect, IL), *10* μ*l protease per 1 ml lysis buffer*, at a ratio of 20 µl lysis buffer/ oocyte. The homogenization was performed by hand until no particles (dark granules) were visible. The homogenized oocytes were then centrifuge at 1,000□×□g, 4°C for 10 minutes to remove cell debris. Next, the supernatant was transferred to a clean centrifuge tube, while the pellet was resuspended with another 20 µl lysis buffer per oocyte. And the centrifugation was repeated. The second supernatant was combined with the first one, and subsequently, this mixture was spun at 200,000□×□g, 4 °C for 45 minutes to obtain oocyte plasma membranes. After the ultracentrifugation, the membrane pellet was resuspended and washed with lysis buffer containing 1 M NaCl at a ratio of 20 µl buffer per oocyte. This mixture was then spun at 200,000□×□g, 4 °C for 45 minutes to collect the membrane pellet. The resulting pellet was further resuspended and washed in lysis buffer containing 1 M NaCl and 2 mM MgCl_2_ at a ratio of 20 µl buffer per oocyte and pelleted down again at 200,000□×□g, 4 °C for 45 minutes.

Afterwards, membrane proteins including RIC-3 were solubilized in solubilization buffer (*1*.*5% Triton X-100, 100 mM K-acetate, 40 mM KCl, 1 mM EDTA, 10 mM MOPS, 0*.*02% NaN3, 2 mM NEM; pH 7*.*5)* supplemented with the protease inhibitor cocktail III, *10* μ*l protease per 1 ml lysis buffer*, at a ratio of 10 μl buffer per oocyte. The solubilization was carried out for 2 hours at 4°C with gently mixing by inverting. After the solubilization was complete, the mixture was centrifuged at 30,000□×□*g*, 4 °C for 1 hour. The supernatant was collected, quickly frozen in liquid nitrogen, and stored at -80°C until use.

### Intracellular domain peptides of 5-HT_3_ receptors

L1-MX and MX peptides of the intracellular domain (ICD) from 5-HT_3_ receptors, Fig. 2A, were designed to contain a C-terminal or N-terminal Cysteine (Cys) with a free sulfhydryl (-SH) group. The peptides were synthesized by GenScript or Biomatik and dissolved at a concentration of 2 mg/ml in ice-cold coupling buffer (50 mM Tris, 5 mM EDTA-Na) or DMSO. Tris (2-carboxyethyl) phosphine (TCEP) was added to the solubilized peptides at a final concentration of 2 to 2.5 mM to prevent oxidation of sulfhydryl groups and maintain sulfhydryls. In case of L1-MX L353A and MX peptides, ice-cold coupling buffer at pH 8.5 and pH 9.5, respectively, was used for initial dissolution; the pH of the peptide solutions was then adjusted to 8.35 and 9.02, respectively, to completely dissolve peptides.

Each peptide concentration was then determined using a nanodrop device (Nanodrop One^C^, ThermoFisher Scientific) by measuring the absorbance at a wavelength of 280 nm or 205 nm^63^, and converted into the respective peptide concentration by using the peptide molecular weight, and peptide extinction coefficient using the following equation:

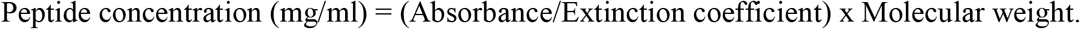

### Pull-down assay of L1-MX peptides and RIC-3

#### Coupling of L1-MX peptides with iodoacetyl resin

Thirty microliters of settled UltraLink™ iodoacetyl resin (ThermoFisher Scientific, # 53155) were added to a Pierce spin column (Thermo Scientific, #69705) and allowed to settle for 5 minutes at room temperature. The liquid was drained by spinning down at 1,000 g for 10 seconds. The resin was then washed three times with 300 µl of coupling buffer (50 mM Tris, 5 mM EDTA-Na, pH 8.5). After removing the buffer of the third wash, peptide with a N- or C-terminal Cys was added to 30 µl settled resin so that a ratio of resin-bound peptide/resin was between 1.72 mg/ml to 34 mg/ml. This specific ratio was determined and optimized for each peptide. To determine the resin-bound peptide/resin ratio, L1-MX peptides were allowed to react with iodoacetyl resin to form thioether bond between the sulfhydryl group of the peptide and the iodacetyl chains of the resin. The coupling reaction was conducted at room temperature for 2 to 5 hours depending on each peptide with gently inverting the spin columns for a few seconds every 30 minutes. Buffer containing unlinked peptide was removed from the spin column by spinning at 1,000 g for 10 seconds. The protein concentration in the flow-through was then determined with the NanoDrop One using the absorbance at 280 nm or 205 nm^63^, and peptide concentrations determined as described above. The unlinked peptide mass was calculated from the peptide concentration and the flow through volume. This mass represents unbound peptide and was subtracted with the peptide mass added to the spin column before the coupling reaction to obtain the resin-bound peptide mass. This mass was then divided by 30 µl resin to obtain the ratio of resin-bound peptide/resin.

After the coupling, resin-bound peptides were washed with 300 µl of the coupling buffer, pH 8.5 for 3 times, and available free iodoacetyl sites of the resin were blocked using 120 µl of freshly made 50 mM L-Cysteine containing 25 mM TCEP, pH 8.5. For control pull-down samples, resin samples without coupled peptide were prepared similarly by modifying the iodoacetyl sidechains with Cys. The blocking reaction was conducted at room temperature with gently inverting spin columns for a few seconds every 10 minutes for 2 hours. The liquid was then removed by spinning at 1,000 g for 10 seconds, and the resin-bound peptide was washed separately with 600 µl of the coupling buffer, 600 µl of 1M NaCl, and 600 µl of storage PBS buffer (0.1 M Na-phosphate, 0.15 M NaCl, 0.05 % NaN_3_, pH 7.2). The peptide was then stored in the PBS buffer at 4°C until use.

#### Pull-down assay with RIC-3

Resin-bound peptides were prepared as described above with a minimum ratio of peptide/resin of 1.72 mg/ml. Storage PBS buffer was removed from the peptides by spinning at 1,000 g for 10 seconds. The resin-bound peptides were then washed 3 times with 300 µl of binding buffer (*0*.*5% Triton X-100, 100 mM K-acetate, 40 mM KCl, 1 mM EDTA, 10 mM MOPS, 0*.*02% NaN*_*3*_; *pH7*.*5)*. The binding buffer was removed by spinning at 1,000 g for 10 seconds. RIC-3 solubilized in 1.5% Triton X-100 was prepared as described above. The protein was thawed on ice for 20 minutes and then centrifuged at 30,000 g for 1 hour at 4°C before incubating with the peptide-resins. The supernatant was collected, and 40 µl of this sample was added to 30 μl of either resin alone or peptide-bound resin. The mixture was incubated at 4°C for 2 hours. Afterward, 150 µl of washing buffer 1 (*1*.*2% Triton X-100, 100 mM K-acetate, 40 mM KCl, 1 mM EDTA, 10 mM MOPS, 0*.*02% NaN3; pH 7*.*5*) was slowly applied to the wall of the spin column without disturbing the resin. The buffer was left in the column for 5 minutes before being removed at 1,000 g for 10 seconds at room temperature (RT). In the next step, 60 µl of washing buffer 2 (*1M NaCl, 25 mM Tris-Cl, 1mM EDTA; pH 7*.*5*) was slowly applied to the wall of the spin column without disturbing the resin. The buffer 2 was drained by spinning at 1,000 g for 10 seconds at RT.

After washing, RIC-3 protein was eluted from resin-bound peptides by applying 40 µL of elution buffer (*277*.*8 mM Tris-HCl, pH 6*.*8, 44*.*4% glycerol, 10% SDS, 10% 2-mercaptoethanol and 0*.*02% Bromophenol Blue*) and incubating at 75°C for 5 minutes, before spinning down at 8,000 g for 3 minutes at RT. The flow through was analyzed using western blot /immunoblot as described below to detect the absence or presence of RIC-3.

### Immunoblot analysis

The entire volume of each flow through obtained from the pull-down assay was separated on Mini-PROTEAN TGX gels (Bio-Rad) and proteins transferred to polyvinylidene fluoride (PVDF) membranes (Bio-Rad) using the Bio-Rad trans-blot turbo system following the manufacturer’s protocol. PVDF membranes were briefly immersed in methanol and then equilibrated in western transfer buffer (20% ethanol, 20% of Bio-Rad 5x transfer buffer, #10026938) for 3-5 minutes. Proteins were then transferred to PVDF membranes at 2.5 Amperes and up 25 Volts for 4 minutes at room temperature. PVDF membranes were blocked under agitation in 5% nonfat milk in tween-containing tris-buffered saline buffer (*TTBS buffer: 0*.*1% Tween-20, 100 mM Tris, 0*.*9% NaCl, pH 7*.*5*) for one hour. Afterwards, the blot was incubated with RIC-3 primary/mouse antibody (Abnova Corporation, Cat # H00079608-B01P) in a dilution of 1:2,000 overnight at 4°C with gentle shaking. Each blot was then washed 3 times, each for 5 minutes in TTBS before incubation with goat raised anti mouse secondary antibody, conjugated to horseradish peroxidase enzyme, HRP (ThermoFisher Scientific product # 31430), in a dilution of 1:5,000 for two hours, under gentle agitation. Subsequently, the blot was washed five times, each for 5 minutes with 20 ml of TTBS buffer. After an additional wash with tris-buffered saline (100 mM Tris, 0.9% NaCl, pH 7.5), SuperSignal West Femto Maximum Sensitivity Substrate (Thermo Scientific) was used for visualizing RIC-3 signal with a digital imaging system (ChemiDoc MP, Bio-Rad).

### Electrophysiology – *Xenopus laevis* oocyte interaction assay

Changes in the functional surface expression of 5-HT_3A_-wt receptor and Ala substitution constructs in the presence and absence of RIC-3 protein were probed using the two-electrode voltage-clamp (TEVC) technique three days post-injection of oocytes. TEVC data were acquired using a TEV-200A Voltage Clamp amplifier (Dagan Corporation), Digidata 1440A digitizer (Molecular Devices), and Clampex 10.7 with gap-free acquisition mode and 200 Hz sampling rate. Resistances of recording electrodes were in the range of 1-5 MΩ. The ground electrode was made of 1% agarose in 3M KCl and connected to a bath and a 200 μl perfusion chamber containing Ringer’s buffer or the OR2 buffer *(82*.*5 mM NaCl, 1 mM MgCl, 20 mM KCl, 5 mM HEPES; pH 7*.*5)*. The voltage and current electrodes were filled with 3M KCl solution. Oocytes were voltage-clamped at a holding potential of -60 mV and continuously superfused under gravity application at 2-5 ml/minute. Currents of 5-HT_3A_ receptors and engineered constructs expressed at the cell surface were evoked by applying 10 μM serotonin creatinine sulfate complex (Sigma), or in short 5-HT, in OR2 buffer. Current amplitudes were obtained by subtracting the maximum current in the presence of 5-HT to the baseline or resting current in the absence of 5-HT. Currents recorded from oocytes injected with cRNA for both RIC-3 and 5-HT_3A_ were recorded and analyzed similarly using Clampfit 10.7. Data were analyzed using GraphPad Prism 6 software and the statistical significance was determined using one-way ANOVA and Tukey’s post-hoc test.

## Abbreviations

(5-HT_3_): The serotonin receptor type 3
(nAChR): nicotinic acetylcholine receptor
(pLGICs): pentameric ligand-gated ion channels
(PPI): protein-protein interaction

## Acknowledgments

We thank the TTUHSC Core Facilities; some of the images and/or data were generated in the Image Analysis Core Facility and Molecular Biology Core Facility supported by TTUHSC.

## Funding and additional information

Research reported in this publication was supported in part by the National Institute of Neurological Disorders and Stroke of the National Institutes of Health under award number R01/R56NS077114 (to M.J.). The funders had no role in study design, data collection and analysis, decision to publish, or preparation of the manuscript.

